# Prevention of EAE by PEGylated Antigenic Peptides

**DOI:** 10.1101/2020.09.08.280875

**Authors:** Jennifer Pfeil, Mario Simonetti, Uta Lauer, Bianca von Thülen, Pawel Durek, Christina Poulsen, Justyna Pawlowska, Matthias Kröger, Ralf Krähmer, Frank Leenders, Ute Hoffmann, Alf Hamann

**Author notes:** Correspondence to: Alf Hamann, Deutsches Rheuma-Forschungszentrum Berlin, Experimental Rheumatology, Charitéplatz 1, 10117 Berlin, Germany. Joint senior authorship (Ute Hoffmann, Alf Hamann).

## Abstract

The treatment of autoimmune disorders such as multiple sclerosis (MS) so far relies largely on the use of non-specific immunosuppressive drugs, which are not able to cure the disease. Presently, approaches to induce antigen-specific tolerance e.g. by peptide-based tolerogenic “inverse” vaccines regain interest. We previously have shown that coupling of peptides to carriers can enhance their capacity to induce regulatory T cells *in vivo*. We here investigated in an experimental autoimmune encephalomyelitis (EAE) model for chronic MS (MOG C57BL/6) whether the tolerogenic potential of immunodominant myelin T cell epitopes can be improved by conjugation to the synthetic carrier polyethylene glycol (PEG). Indeed, preventive administration of the PEGylated antigenic peptide could almost completely protect mice from EAE development, which was accompanied by reduced immune cell infiltration in the central nervous system (CNS). Depletion of Tregs abrogated the protective effect indicating that Tregs play a crucial role in induction of antigen-specific tolerance in EAE. Treatment during the acute phase was safe and did not induce immune activation. However, treatment at the peak of disease was not affecting the disease course, suggesting that either induction of Tregs is not occurring in the highly inflamed situation, or that the immune system is refractory to regulation in this condition. Thus, PEGylation of antigenic peptides is an effective and feasible strategy to improve tolerogenic (Treg-inducing) peptide-based vaccines, but application in overt disease might require modifications or combination therapies that simultaneously suppress effector mechanisms.

## 1 Introduction

Chronic autoimmune and inflammatory diseases are affecting an increasing number of individuals. Current treatments, including biologics, are not effective in all patients, do not provide cure of the disease and might be associated with severe side effects. This continuing medical need has led to regained interest in antigen-specific tolerogenic therapies, also termed “inverse vaccination” (Dolgin, 2010; Willekens and Cools, 2018). The phenomenon of immunological tolerance is known since the beginnings of cellular immunology and partially been used in the specific immunotherapy (SIT) in allergy. However, its application to autoimmune diseases is still awaiting major breakthroughs, despite the obvious advantage of treating selectively pathogenic mechanisms instead of applying unspecific immunosuppressants.

For several autoimmune diseases, including MS, major autoantigens and their immunodominant domains are known and animal models for MS have provided strong evidence for a role of antigen-specific T cells in the pathogenesis (Martin et al., 2016). This is the rationale for approaches to generate peptide-based tolerogenic vaccines for the treatment of MS.

Initially, intravenous administration of protein, immunodominant peptides or altered peptide ligands with a varied amino acid sequence were found to be tolerogenic and to induce Tregs in experimental models (Anderton et al., 1999; Sospedra and Martin, 2005; Miller et al., 2007). However, clinical trials were not effective so far, and in some preclinical or clinical studies adverse effects occurred (Genain et al., 1996; Bielekova et al., 2000; Kappos et al., 2000; Smith et al., 2005). Obstacles in the use of peptides could comprise their short half-life in serum due to rapid excretion through the kidney or enzymatic degradation, or unfavorable physicochemical properties (aggregation) of some peptides probably being a reason for anaphylactic reactions. Further improvements therefore included the loading of apoptotic cells or of synthetic carriers with antigenic peptides, use of tolerogenic adjuvants, or a combination thereof (Pearson et al., 2017; Ben-Akiva et al., 2018; Kishimoto and Maldonado, 2018; Feng et al., 2019; Pearson et al., 2019).).

Approaches of peptide-modified autologous cells (Smith and Miller, 2006) entered already the stage of clinical trials (Lutterotti et al., 2013; Zubizarreta et al., 2019). However, cell-based medicinal products are poorly defined and require an enormous personalized production process as well as extensive regulatory requirements. While micro- and nanoparticles avoid some of these drawbacks and showed promising results, they still bear significant difficulties regarding standardization and approval.

We therefore considered that a more simple chemical modification could lead to peptide-based vaccines with improved tolerogenic activity. We recently reported that peptides conjugated to oligo- or polyglycerols or to a repetitive protein are superior in the induction of regulatory T cells compared to native peptides (Gupta et al., 2015; Puentes et al., 2016). These findings were compatible with the view that increasing the size of a peptide by coupling to a macromolecular carrier of any kind would result in a better tolerogenic effect either by increasing the short half-life of native peptides in the circulation and/or by improved uptake/storage in antigen-presenting cells (APC).

We therefore investigated whether coupling of myelin peptides to a soluble carrier entity consisting of polyethylene glycol (PEG) would result in increased tolerogenicity. In a preceding paper, we described that indeed, vaccination with PEG-coupled OVA-peptide in the DO11.10 experimental model results in a marked increase in Treg numbers and downregulation of effector T cells (Pfeil et al., Submitted).

PEG is a synthetic polymer composed of repetitive ethylene oxide subunits, either in linear form or as branched polymers. Numerous PEGylated biomolecules have already been approved by the Food Drug Administration (FDA) and the European Medicines Agency (EMA) for human use as ingredients of foods, cosmetics and pharmaceuticals including topical, parenteral and nasal formulations. The covalent attachment of PEG (PEGylation) improves the pharmacokinetic and pharmacodynamic profile of biomolecules by increasing serum half-life as well as proteolytic resistance, while reducing renal clearance and immunogenicity (Greenwald et al., 2003; Fishburn, 2008; Zheng et al., 2012; Mu et al., 2013). We here used conjugates with PEG20 (i.e. consisting of 20 ethylene units) if not otherwise mentioned, which was determined as the most effective carrier type in our preceding study (Pfeil et al., Submitted).

The tolerogenic potential of PEG-coupled myelin peptides were investigated in a murine model for human multiple sclerosis (MS), experimental autoimmune encephalomyelitis (EAE). In the C57BL/6 mouse model largely used here, EAE is induced by the encephalitogenic peptide MOG_35-55_ of myelin oligodendrocyte glycoprotein in complete Freund’s adjuvant (CFA). In this model, EAE displays typically a chronic clinical course (Mendel et al., 1995).

The results show that vaccination with PEGylated myelin peptide exerts a tolerogenic effect that results in protection from EAE development in the MOG C57BL/6 model. However, when applied during the peak of inflammation, a suppression of ongoing disease was not achieved in this active EAE model. This suggests that the PEGylation alone is not sufficient for highly effective therapeutics, however, might be applied during remission or in combination with treatments more efficiently suppressing the effector phase. Treatment with PEGylated peptide was safe, as no adverse immune hyperactivation upon peptide delivery during the peak of disease was observed.

## 2 Material and Methods

### Mice

C57BL/6 mice were either purchased from Charles River or Janvier. Mice were maintained under specific pathogen-free conditions according to national and institutional guidelines in the breeding facility of the Deutsches Rheuma-Forschungszentrum Berlin (DRFZ). All experiments were approved by the Landesamt für Gesundheit und Soziales (LAGeSo).

### Peptides

MOG-peptide 35-55 (pMOG): MEVGWYRSPFSRVVHLYRNGK and the cysteine-modified pMOG: CβA-MEVGWYRSPFSRVVHLYRNGK, used for coupling, were either synthesized in house (Institute for Medical Immunology, Charité Universitätsmedizin Berlin, Germany) or purchased from Pepceuticals (Leicestershire, UK).

### Synthesis of pMOG-PEG20

To couple PEG to the C-modified N-terminus of pMOG, 157 mg of 20 kDa PEG-maleimide (NOF Corporation, Japan) and 20 mg pMOG were reacted in 20 mL of a 20 mmol/l sodium phosphate buffer pH 7.2 containing 250 µmol/l of Tris(2-carboxyethyl)phosphine. The reaction mixture was stirred at RT for 1 h. Progress of the reaction was monitored by RP-HPLC.

### Purification of pMOG-PEG20

Crude peptide-PEG-conjugates were purified by cation-exchange chromatography using MacroCap SP on an Äkta chromatography system. pMOG-PEG20 was bound to the resin in 20 mmol/L sodium acetate pH 4.5. Conjugate eluted with a linear sodium chloride gradient from 0 to 500 mM in 10 column volumes in the same eluent. Conjugate elution was monitored at 213 nm. Fractions containing the peptide-PEG-conjugate was pooled and desalted by dialysis against deionized water. Subsequently, pMOG-PEG20 was concentrated by freeze-drying. Prior to use, conjugate was reconstituted in WFI and filtered through a sterile 0.2 µm filter.

### Analysis of pMOG-PEG20

Analysis of pMOG-PEG20 was performed by MALDI-ToF-MS, UV-spectrometry, and reversed phase-high performance liquid chromatography (RP-HPLC). RP-HPLC was performed on a Waters Alliance System. For the analysis of pMOG-PEG20, a Butyl C4 column 4.6 × 250 mm (5 µm) was used. Sample was eluted with a linear gradient from 70:15:15 to 50:25:25 water:acetonitrile:isopropanol each with 0.1% TFA in 10 min. PEG or peptide-PEG-conjugate were detected using an evaporating light scattering detector (ELSD). The peptide-PEG-conjugates were free of any residual unmodified peptide

### Cell preparation

Single cell suspensions were prepared from spleen by mechanical dissociation, lysis of red blood cells in lysis buffer (0.01 M potassium bicarbonate, 0.155 M ammonium chloride and 0.1 mM ethylenediaminotetraacetic acid (EDTA)) and washed with phosphate buffered saline (PBS) supplemented with 0.2% bovine serum albumin (BSA).

To isolate hematopoietic cells from spinal cord, mice were perfused after sacrifice with 30 ml cold PBS. Spinal cords were minced and digested at 37°C for 15 min in serum free RPMI containing 1 mg/ml collagenase IV (Sigma-Aldrich, Taufkirchen, Germany) and 0.5 mg/ml Dnase I (Sigma-Aldrich). After digestion, spinal cord was flushed through a 18G needle and incubated for a further 15 min at 37°C. After the 2^nd^ incubation the preparation was flushed through a 25G needle to obtain a single cell suspension, which was forced through a 40 µm cell strainer. Hematopoietic cells including microglia were isolated from discontinuous 30%:70% percoll gradients by centrifugation at 2000 rpm for 20 min at RT and stained with the respective marker combinations given below.

### Antibodies and flow cytometry

The following monoclonal antibodies (mAbs) and appropriate isotype controls were obtained from eBioscience (San Diego, CA, USA): eFluor 450-conjugated anti-CD4 (RM4-5), PE-Cy7-conjugated anti-CD11b (M1/70), FITC-conjugated anti-CD45 (30F11), PE-conjugated anti-GM-CSF (MP1-22E9), PE-Cy7-conjugated anti-IFN-γ (XMG1.2), FITC-conjugated anti-IL-17A (eBio17B7), PerCP-eFluor 710-conjugated anti-TNFα (MP6-XT22). Pacific Blue-conjugated anti-CD3 (17A2), PerCP-conjugated anti-CD11c (N418) and Pacific-Blue-conjugated anti-CD19 (6D5) antibodies were obtained from Biolegend (San Diego, USA). The APC-conjugated anti-CD154 antibody was obtained from Miltenyi Biotec. PE-conjugated anti-GR-1 (RB6-8C5), anti-Fcγ-receptor (2.4G2) and anti-CD25 (PC61) antibodies were produced in house (DRFZ). Total rat IgG was purchased from Dianova (Hamburg, Germany).

To stain surface molecules, cells were pre-incubated with anti-Fcγ-Receptor antibody (20 µg/ml) before staining with specific mAbs.

To analyze cytokine production of splenic MOG-specific CD4^+^ T cells in EAE, splenocytes were harvested and 1x 10^7^cells were re-stimulated in presence of 100 µg/ml pMOG in a 96 well plate for 14 h. After 2 h incubation Brefeldin A was added. Cells were surface-stained with an anti-CD4 antibody. After fixation by incubation with 2% paraformaldehyde (PFA, Sigma-Aldrich) at RT, intracellular staining was performed by 5 min pre-incubation with rat IgG in 0.5% saponin (Sigma-Aldrich) and addition of anti-CD154 antibody and anti-cytokine antibodies for another 30 min in 0.5% saponin at RT.

Quantification of hematopoietic cells in the spinal cord was done after isolation by flow cytometric analysis of the following marker using a sequential gating strategy (see Supplementary Figure S4xx): microglia (CD45^int^ CD11b^+^), neutrophils (CD45^high^ GR1^high^ CD19^-^), CD11b^-^ DCs (CD45^high^ GR1^-^ CD11c^+^ CD11b^-^ CD19^-^), CD11b^+^ DCs (CD45^high^ GR1^-^ CD11c^+^ CD11b^+^ CD19^-^), macrophages (CD45^high^ GR1^-^ CD11c^-^ CD11b^+^ CD19^-^), B cells (CD45^high^ CD19^+^) and T cells (CD45^high^ GR1^-^ CD11c^-^ CD11b^-^ CD19^-^ CD3^+^).

### Induction and clinical evaluation of MOG-induced EAE model and treatment protocol

For induction of EAE, female C57BL/6 mice (age, 8–10 weeks) were immunized *s*.*c*. at 4 spots on the flanks with 200 µl of an emulsion of MOG-peptide 35-55 (pMOG, 250 µg per mouse) in complete Freund’s adjuvant (CFA, Difco, Heidelberg, Germany) supplemented with 800 µg of *Mycobacterium tuberculosis* H37Ra on day 0. Mice also received *i*.*p*. 200-400 ng of *Bordetella pertussis* toxin (List Biological Laboratories, Campbell, USA) on day 0 and 2. Paralyzed mice were given easier access to food and water.

#### Prevention approach

C57BL/6 mice were tolerized *i*.*v*. with PBS (control), 7.6 µg pMOG or equimolar amounts of pMOG-PEG20 7, 14 or 28 days prior to EAE induction.

#### Treatment approach

C57BL/6 mice received *i*.*v*. PBS (control), 7.6 µg pMOG or equimolar amounts of pMOG-PEG20 7 days post EAE induction. Pilot experiments using the relapsing-remitting MBP-PLP model in SJLxB10.PL mice (Miller et al., 2010) were carried out as described in Supplementary Materials.

Individual animals were monitored every day, and clinical scores were assessed as an accumulative score as follows: TPA tail paresis 0.5; TPA-L almost complete tail plegia 0.75; TPL tail plegia 1; (RRW) righting reflex impaired 0.25; RRW righting reflex weak 0.5; HPA hind limb paresis 0.5; HPA-L almost complete hind limb plegia 1; HPL hind limb plegia 1.5; FPA fore limb paresis 1.

For Treg depletion, animals were treated *i*.*p*. with 500 µg anti-CD25 antibody 7 days prior to EAE induction.

### Statistics

Significance was determined with the non-parametric Mann-Whitney U test using PRISM 5.02 (GraphPad, La Jolla, CA, USA). Differences were considered as statistically significant with p≤0.05 (*), very significant with p≤0.01 (**) and extremely significant with p≤0.001 (***).

## 3 Results

### Preventive administration of pMOG-PEG20 suppresses chronic experimental autoimmune encephalomyelitis

The tolerogenic effect of systemic administration of PEGylated MOG-peptide was tested in the C57BL/6 EAE mouse model. Mice received PBS (control), 7.6 µg encephalitogenic pMOG_35-55_ (equivalent to 14.7 µM; 0.38 mg/kg) or equimolar amounts (based on peptide amount) of PEG20 conjugated to the encephalitogenic pMOG_35-55_ 7 days prior to EAE induction. In contrast to pMOG, which displayed only a weak tolerogenic effect, pMOG-PEG20 pretreatment almost completely suppressed disease development (Figure 1).

**Figure 1.**
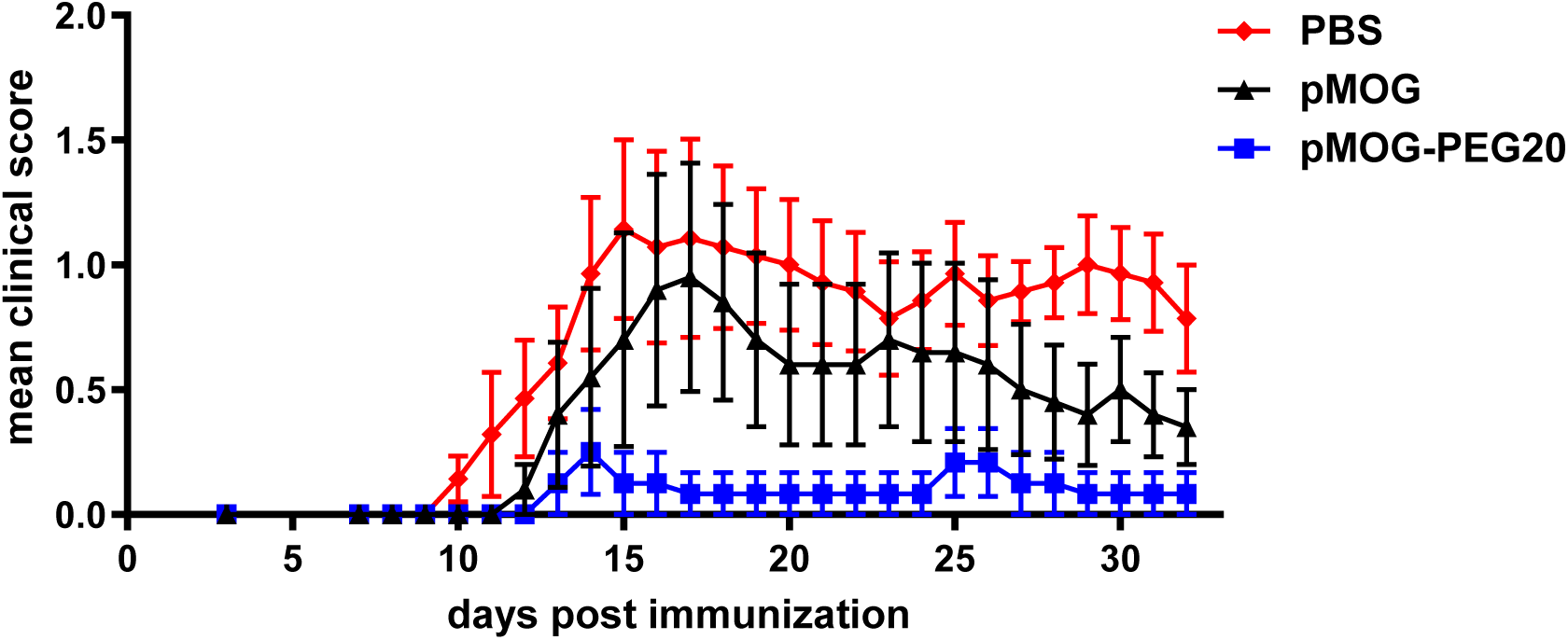
Improved tolerogenicity of pMOG-PEG20 compared to unconjugated peptide in EAE. EAE was induced by *s*.*c*. immunization of C57BL/6 mice with pMOG_35-55_ in complete Freund’s adjuvans (CFA) containing *Mycobacterium tuberculosis* H37Ra on day 0. In addition, mice received *Bordetella pertussis i*.*p*. on day 0 and day 2. 7 days prior to EAE induction mice were tolerized with PBS (control), 7.6 µg pMOG or equimolar amounts (based on peptide amount) of pMOG-PEG20. Individual animals were observed every day, and clinical scores were assessed as an accumulative score. Mean clinical score per group ± SEM. PBS n = 7; pMOG n = 5; pMOG-PEG20 n = 6. One representative of 3 independent experiments is shown.

Pretreatment with pMOG conjugated to a higher molecular weight PEG, PEG40, was as efficient as pMOG-PEG20 vaccination (Supplementary Figure S1).

Tolerization with either pMOG or pMOG-PEG20 was less effective when done 4 weeks instead of 1 week before disease induction (Supplementary Figure S2); tolerization at 2 weeks before induction was in 3 out of 4 experiments as complete as in the 1-week setting (data not shown). These data demonstrate that vaccination with PEGylated peptide is able to prevent the development of EAE, although a single tolerogenic vaccination might not be sufficient to induce a long-term tolerant state.

### Tregs play a role in antigen-specific tolerance induction by pMOG-PEG20

In the DO11.10 adoptive transfer model, we demonstrated that vaccination with PEGylated peptides led to expansion and/or *de novo* induction of antigen-specific Tregs (Pfeil et al., Submitted). To analyze the role of antigen-specific Tregs in the prevention of EAE by pMOG-PEG20 pretreatment, tolerogenic vaccination was given 2 weeks prior to EAE induction and Tregs were depleted by injection of anti-CD25 antibody 7 days prior to EAE induction. As expected, Treg depletion enhanced disease severity (Figure 2A, B).

**Figure 2.**
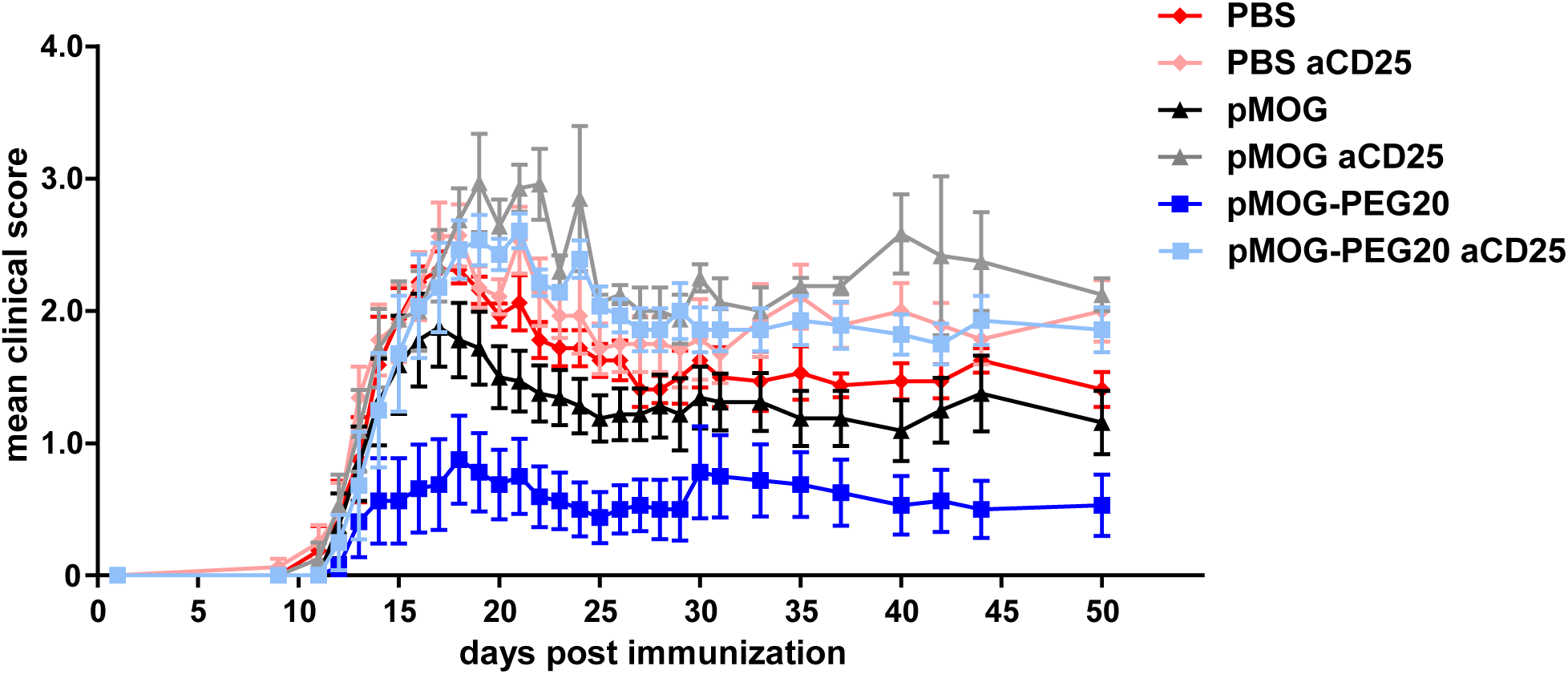
Tregs are involved in protective tolerance induced by pMOG-PEG20. 14 days prior to EAE induction, C57BL/6 mice were tolerized with PBS (control), 7.6 µg pMOG or equimolar amounts of pMOG-PEG20. After 7 days mice received either 500µg anti-CD25 (PC61) antibody or PBS (control) *i*.*p*.. EAE was induced as described above. Mean clinical score per group ± SEM, n= 7-8 per group. One representative of 4 independent experiments is shown.

These findings suggest that Tregs play a crucial role in EAE suppression induced by tolerogenic pMOG-PEG20 vaccination.

### Effects on the frequency of MOG-specific splenocytes among total CD4^+^ cells and their effector cytokine profile upon prior administration of pMOG-PEG20

In a preceding study (Pfeil et al., Submitted) in the DO11.10 adoptive transfer model we demonstrated that vaccination with PEGylated peptide resulted in partial depletion of specific T cells as well as reduction of pro-inflammatory cytokine producers in spleen. To investigate whether the protective effect in EAE of pMOG-PEG20 is also involving modulation of effector cells, we analyzed the frequency of MOG-specific cells among total CD4^+^ T cells and their pro-inflammatory cytokine profile by FACS analysis of pMOG-restimulated T cells at different time points.

The frequency of MOG-specific cells (CD40L^+^ upon pMOG-restimulation) among total CD4^+^ splenocytes was neither in the pre-onset phase nor at peak of disease changed after previous vaccination with pMOG or pMOG-PEG20. (Figure 3 A-C:) The frequency of TNF- and IFN-γ-producing cells among MOG-specific (CD4^+^ CD40L^+^) splenocytes was significantly decreased by pMOG-PEG20 prior disease onset (d7 post EAE induction), reflecting partial anergy of MOG-specific cells (Figure 4 A, B). Also frequencies of IL-17A- and GM-CSF-producing cells among antigen-specific cells appeared to be reduced in mice tolerized with pMOG-PEG20 compared to non-tolerized mice albeit not reaching significance (Figure 4 C, D). At peak of disease, hardly any significant changes were seen (Supplementary Figure S3). Thus, preventive vaccination in the MOG-model does not result in a depletion of MOG-specific cells and only a weak downregulation or reduction of cytokine producing T cells can be observed.

**Figure 3.**
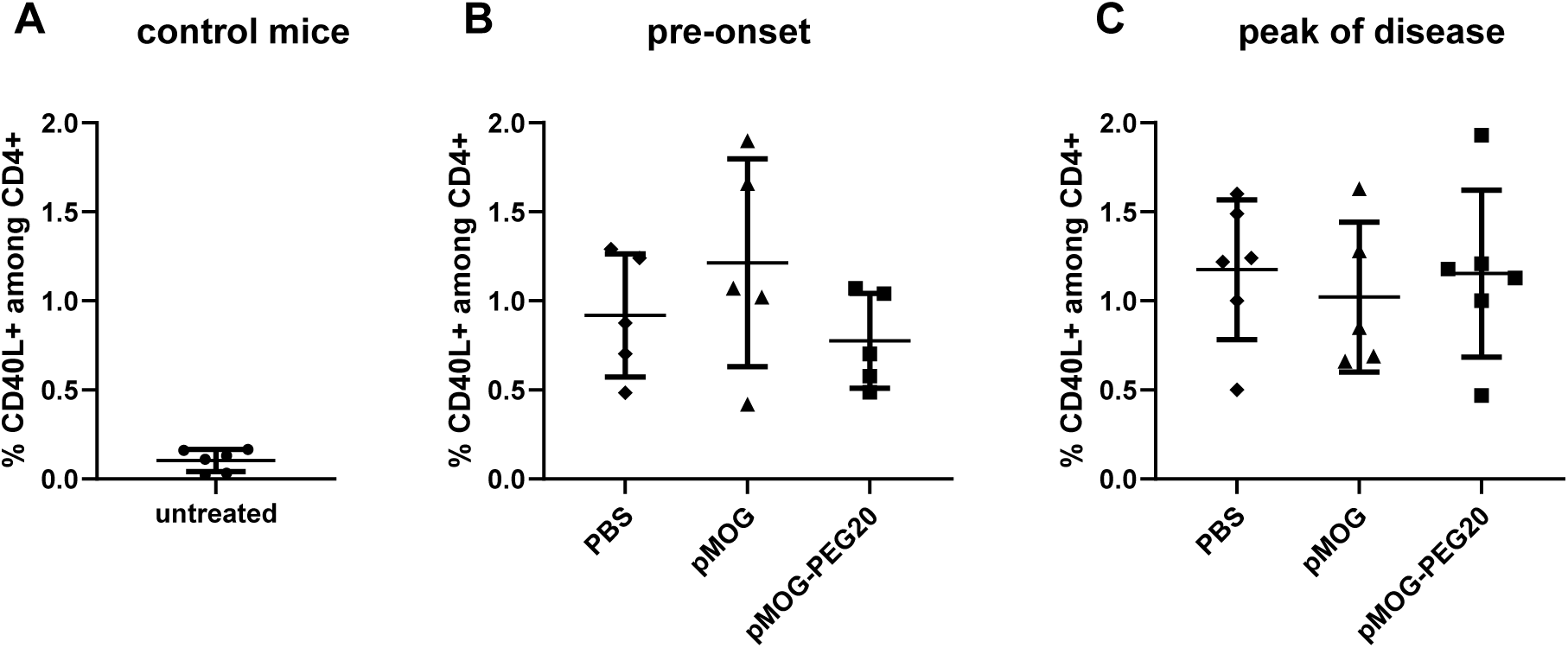
In the pre-onset phase and at the peak of EAE, the frequency of MOG-specific cells among total CD4^+^ splenocytes is not affected by pMOG-PEG20. Splenocytes were re-stimulated overnight with pMOG and analyzed by flow cytometry. Data represent means ± SD of % MOG-specific CD4^+^ cells (% CD40L^+^ upon pMOG-restimulation) among total CD4^+^ cells. (**A**) untreated control mice, n = 6. (**B**) pre-onset phase: 7 days post EAE induction, n = 5. (**C**) peak of disease: 15 days post EAE induction, n = 5. C57BL/6 mice were tolerized 7 days prior to EAE induction *i*.*v*. with either PBS (control), 7.6 µg pMOG or an equimolar amount of pMOG-PEG20 (**B, C**). One representative of 2 independent experiments is shown. P values were determined by unpaired non-parametric Mann-Whitney U test.

**Figure 4.**
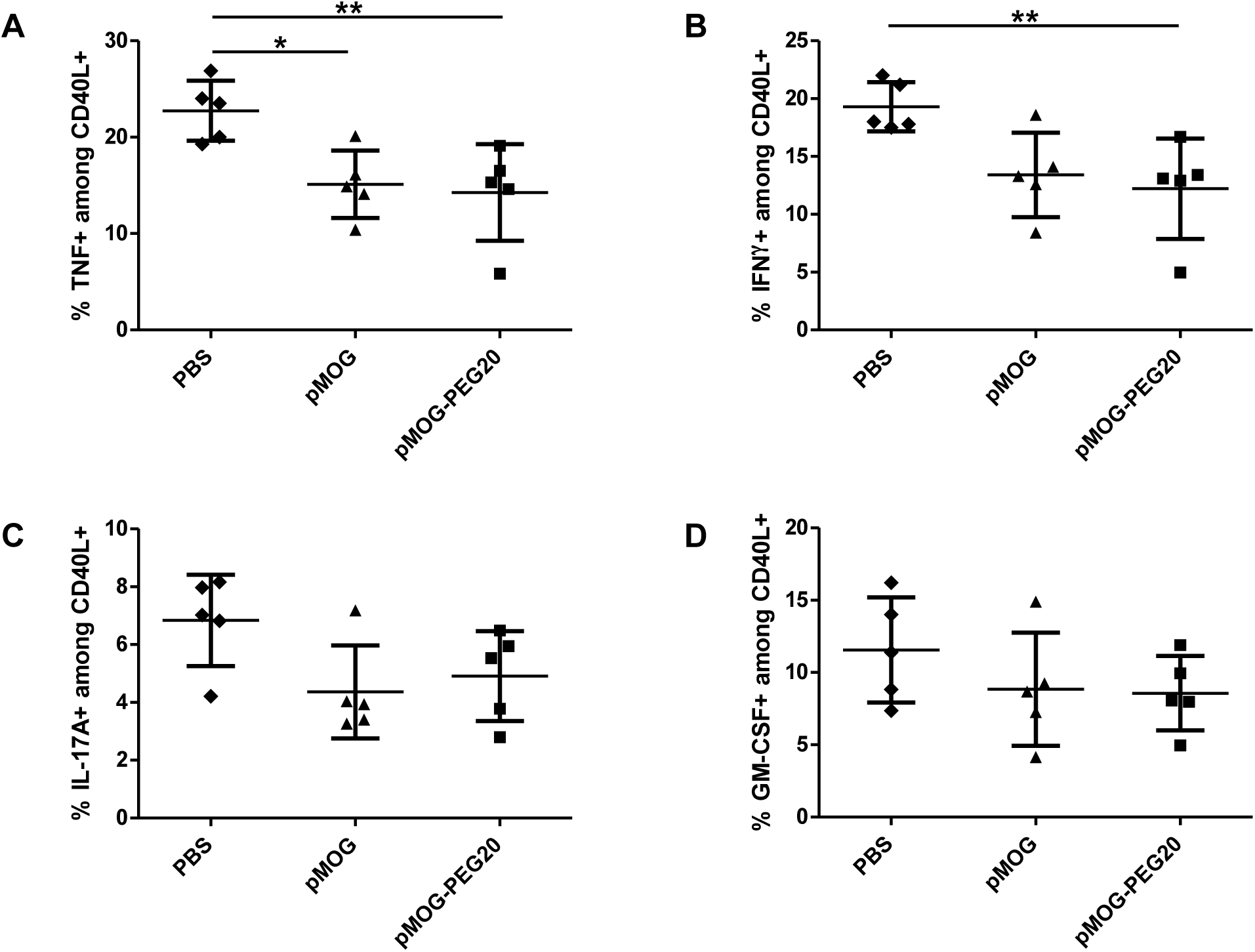
In the pre-onset phase of EAE, the frequency of TNF or IFN-γ-producing cells among MOG-specific splenocytes is significantly decreased compared to PBS by pMOG-PEG20 and, slightly less efficient, by pMOG. 7 days prior to EAE induction, C57BL/6 mice were tolerized *i*.*v*. with either PBS (control), 7.6 µg pMOG or an equimolar amount of pMOG-PEG20. Animals were sacrificed 7 days (pre-onset phase) post EAE induction. Splenocytes were re-stimulated overnight with pMOG and analyzed by flow cytometry. Data represent means of (**A**) % TNF^+^, (**B**) % IFN-γ^+^, (**C**) % IL-17A^+^ and (**D**) % GM-CSF^+^ splenocytes among CD4^+^ CD40L^+^ (MOG-specific) cells ± SD (n = 5 per group). One representative of 2 independent experiments is shown. P values were determined by unpaired non-parametric Mann-Whitney U test.

### Administration of pMOG-PEG20 reduces CNS immune cell infiltration

Cellular infiltration is a hallmark of inflammation. To analyze whether tolerogenic pMOG-PEG20 vaccination reduces the accumulation of inflammatory cells in the CNS, the composition of different immune cell populations in the tissue was analyzed. Spinal cord was isolated from tolerized and control pMOG/CFA-immunized as well as from untreated mice at the peak of the disease. Cellular infiltrates were analyzed after dissociation by flow cytometry. Among viable CD45^+^ cells, microglia, neutrophils, DC subsets, macrophages as well as T and B cells were identified using a sequential gating strategy (Supplementary Figure 4).

pMOG-PEG20 vaccination not only diminished infiltrating T cells but also total leukocytes in the CNS cells compared to pMOG and PBS application, in particular the infiltration by neutrophils, macrophages and CD11b^+^ DCs was strongly reduced (Figure 5; for relative proportions of subsets and statistics see Supplementary Figure S5 A, B). This, in fact, correlates with reduced EAE symptoms following pMOG-PEG20 vaccination.

**Figure 5.**
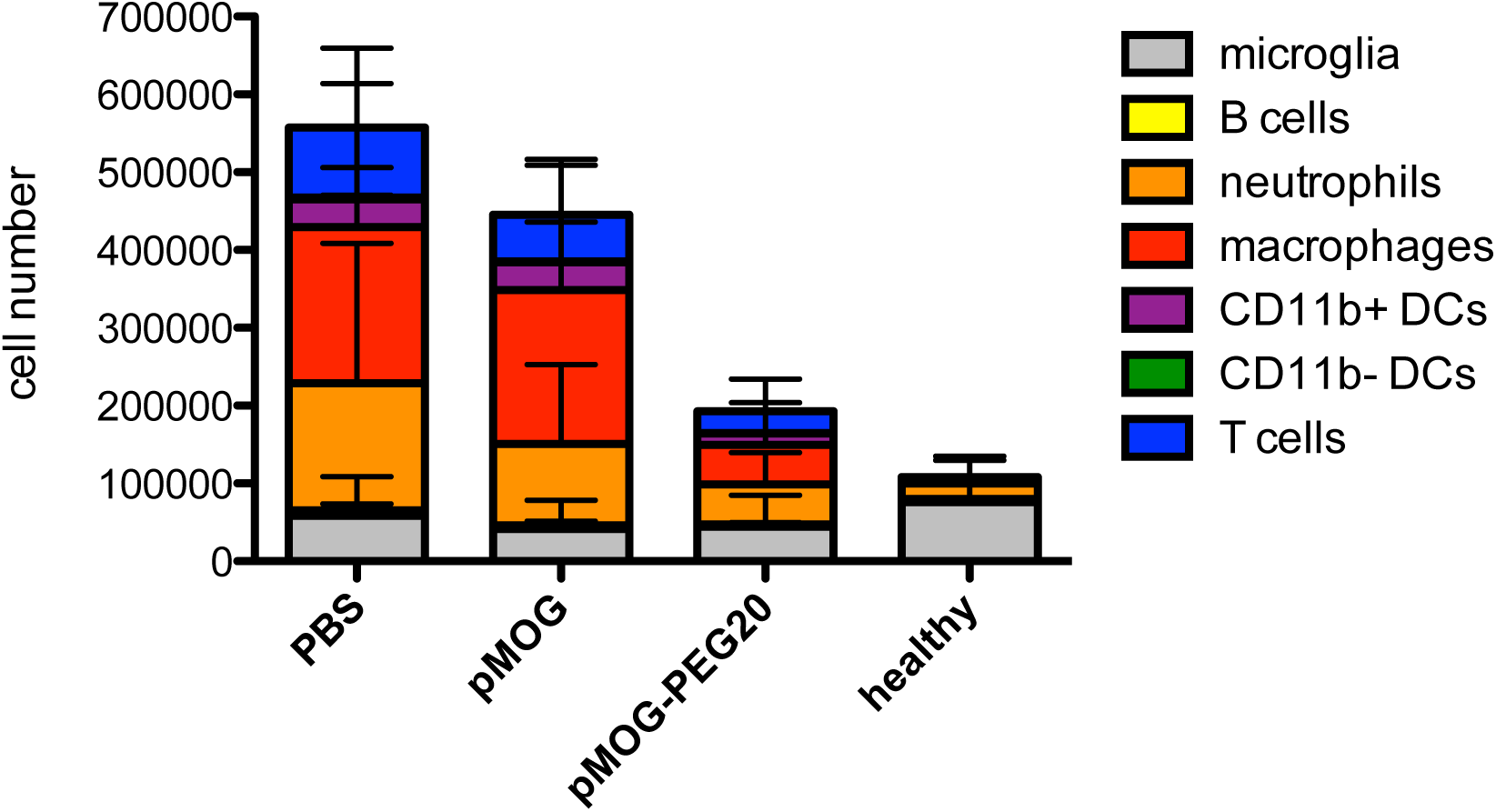
Reduced CNS infiltration at peak of disease following pMOG-PEG20 vaccination. 7 days prior to EAE induction mice were tolerized with PBS, 7.6 µg pMOG or equimolar amounts of pMOG-PEG20. At the peak of disease (day 15) spinal cords were isolated from pMOG-immunized C57BL/6 mice as well as from non-immunized untreated C57BL/6 mice (control). Different leukocyte subsets were quantified by flow cytometric analysis of marker combinations as described in Methods and Supplementary Figure S4. Data are depicted as mean ± SD (n = 6-12). Data is summarized from 2 independent experiments. *P* values were determined by unpaired non-parametric Mann-Whitney U test and are listed in Supplementary Table S5.

### Administration of pMOG-PEG20 in established disease is ineffective

In order to determine the potential application of tolerogenic vaccination with PEGylated peptides in acute phases of autoimmune disease, we investigated whether therapeutic administration of pMOG-PEG20 is effective in ongoing CNS inflammation. Mice received PBS, 7.6 µg pMOG or equimolar amounts (based on peptide amount) of pMOG-PEG20 7 days after EAE induction. Therapeutic administration of both pMOG-PEG20 and pMOG could not suppress EAE symptoms (Figure 6). Of note, treatment with PEGylated pMOG did however not lead to exacerbation of the disease.

**Figure 6.**
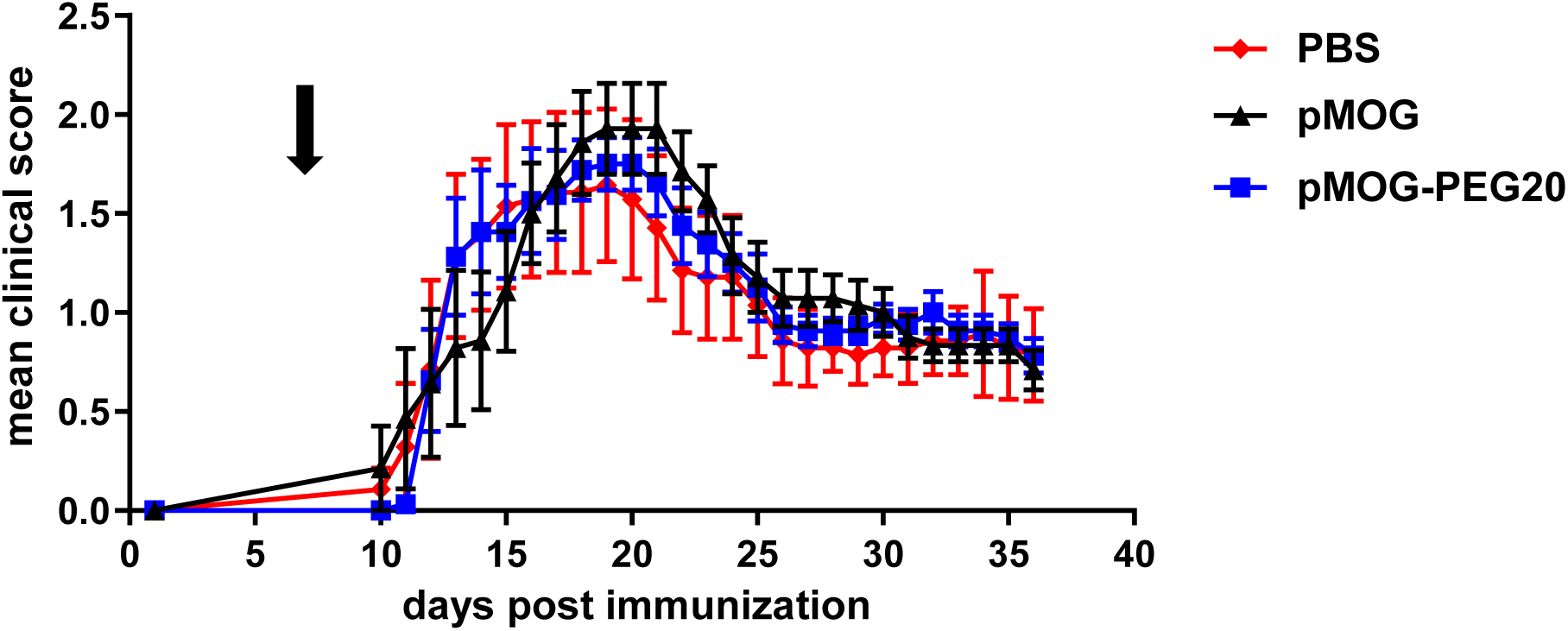
Therapeutic administration of pMOG-PEG20 does not prevent EAE development, but also does not lead to exacerbation of the disease. **7** days post EAE induction, C57BL/6 mice received PBS (control), 7.6 µg pMOG or equimolar amounts of pMOG-PEG20. Mean clinical score per group ± SEM (PBS n = 7; pMOG n = 7; pMOG-PEG20 n = 8). One representative of 2 independent experiments is shown.

These data demonstrate that the inflammatory milieu in ongoing disease prevents successful tolerization with either peptide or PEG-modified peptide under the used conditions.

In pilot experiments, we also applied PEGylated PLP-peptides in the relapsing-remitting MBP-PLP model in SJLxB10.PL mice. Also repeated application of PEGylated PLP-peptide at high doses was not ameliorating disease apart from a delay of the first relapse (Supplementary Figure S6).

## 4. DISCUSSION

Tolerogenic vaccines based on immunodominant peptide sequences from autoantigens are an attractive concept for the development of antigen-specific therapies of autoimmune diseases. While readily convertible into a pharmaceutical product, efficacy in clinical trials was disappointing, giving rise to the development of new approaches. We here investigated whether coupling of myelin peptides to defined and clinically well-proven PEG moieties would increase their tolerogenic potential and therapeutic effect in EAE.

Indeed, EAE development was almost completely suppressed by vaccination with pMOG-PEG20, while free peptide only led to a modest amelioration of EAE. This is in line with our concept that increasing the size of autoantigenic peptides by coupling to an inert carrier is improving their tolerogenic potential.

This concept evolved from previous studies showing protective effects of peptide coupled to repetitive protein domains in EAE (Puentes et al., 2016) and the efficacy of PEG- as well as oligo-or polyglycerol-conjugated peptides in inducing Tregs and reducing T effector cells in a transgenic OVA-model (Gupta et al., 2015; Pfeil et al., Submitted). In the OVA-model we found that size of the PEG carrier was critical, with linear PEG20 (20 kDa) being the most effective inducer of Tregs among 12-40 kDa variants and a tetrameric variant (Pfeil et al., Submitted). We therefore used in the present study predominantly MOG-peptide 35-55 coupled to PEG20 (pMOG-PEG20), although pMOG-PEG40, was hardly less efficient in preventing EAE. We demonstrated previously that PEGylation leads to a major prolongation of bioavailability of the tolerogenic peptides, either caused by the known increase in serum half-life or additional effects such as storage in APC as discussed elsewhere (Pfeil et al., Submitted). It can be assumed that this is also the main mechanism of action in the improved tolerogenicity of the other types of carriers mentioned above.

As compared to the aforementioned different scaffolds and to encapsulation or coupling of peptides into and onto nanobeads (Ben-Akiva et al., 2018; Kishimoto and Maldonado, 2018; Feng et al., 2019; Pearson et al., 2019), PEG has the advantage of being widely used already in the pharmacological optimization of biologicals and is clinically proven, thus synthesis and regulatory processes might bear less hurdles. The clinical application of PEGylated compounds has, however, also uncovered some issues caused by pre-existing antibodies to PEG, most likely originating from the widespread use of PEG in cosmetics etc., (Zhang et al., 2016). While anti-PEG antibodies were not affecting efficacy e.g. in trials using PEGylated interferon, in other cases the biokinetics of PEGylated bacterial enzymes were found to be negatively affected in some patients (Poppenborg et al., 2016; Zhang et al., 2016). Whether the type of conjugates used here is affected by anti-PEG antibodies remains to be observed. It should be mentioned that there are presently no studies available on the potential formation of antibodies to other synthetic carriers or nanobeads.

In our previous studies (Gupta et al., 2015; Puentes et al., 2016; Pfeil et al., Submitted) we demonstrated that coupling of peptides to carriers improved the induction or expansion of antigen-specific Tregs and it became obvious that linker chemistry, size and structure beside dose played a critical role for the Treg-inducing potential. To investigate whether Tregs are critical mediators of the tolerogenic effect also in the EAE model, Tregs were depleted with anti CD25 Ab. Indeed, the protective effect of vaccination with pMOG-PEG20 was completely neutralized upon depletion. These findings confirm previous studies showing that Tregs play a significant role in the regulation of EAE and MS (reviewed in (Anderton and Liblau, 2008; Danikowski et al., 2017). Interestingly, in case of EAE prevention by apoptotic, antigen coupled leukocytes, Tregs were shown to be dispensable for tolerance induction but essential for long-term tolerance maintenance (Getts et al., 2011).

While the prominent role of Tregs in regulating the immunological balance and keeping self-tolerance became clear in the last two decades, previous studies, notably such in high-zone tolerance also pointed to mechanisms such as depletion of antigen-specific cells or induction of anergy or adaptive tolerance, i.e. a state of unresponsiveness including a suppression of (effector-) cytokine production (Choi and Schwartz, 2007). In line with the studies in the OVA-model (Pfeil et al., Submitted), a moderate but significant reduction in the percentage of MOG-specific cells able to produce the effector cytokines TNF or IFN-γ and a non-significant trend for reduction of producers of IL-17A and GM-CSF was observed, albeit only in the pre-onset phase. All these cytokines have been considered as drivers in the pathogenesis of MS (Croxford et al., 2015; Kaskow and Baecher-Allan, 2018; McGinley et al., 2018; Pegoretti et al., 2018). Whether the moderate reduction in cytokine producers observed here contributes to tolerance in the present model remains to be shown. It should be mentioned that MOG-specific cytokine producers could only be determined in cells isolated from the spleen; it is not excluded, that relative numbers are different in the inflamed CNS tissue as effector-memory cells tend to be trapped in sites of inflammation (Ghani et al., 2009; Marelli-Berg et al., 2010). Hence, the present data do not allow a firm conclusion as to the role of this partial anergy induction for the tolerogenic effect of vaccination.

The superior effect of PEGylated pMOG as compared to native pMOG seen in clinical scores was mirrored in the strong reduction of inflammatory cells accumulating in the diseased spinal cords upon vaccination. Although the influx of inflammatory cells was not completely prevented by vaccination with pMOG-PEG20, total hematopoietic cells were significantly reduced to approximately 1/3. Notably macrophages and neutrophils that have both been implicated in pathology of MS (Mishra and Yong, 2016; De Bondt et al., 2020) were strongly reduced. This underlines the protective effects of pMOG-PEG20 vaccination recorded in the clinical symptoms.

With a few exceptions, the intended clinical use of tolerogenic approaches in the treatment of autoimmune diseases or allergy is rather not prevention of disease but treatment of an ongoing autoimmune process. Here, the preceding or continuing activation of the immune system bears the most critical hurdle for efficacy. Both the generation and the efficacy of Tregs have been reported to be impaired under inflammatory conditions (Thorstenson and Khoruts, 2001; Belkaid and Oldenhove, 2008; Caretto et al., 2010). In the OVA-model, we observed that concomitant injection of LPS completely prevented the induction of Tregs upon tolerogenic vaccination with pOVA-PEG20 (Pfeil et al., Submitted).

It was therefore not completely unexpected that application of pMOG-PEG20 (as pMOG) were unable to change the disease course when given after induction of EAE. We reasoned that a remitting-relapsing model might bear windows of opportunity for the induction of Tregs during ongoing disease. However, this was not found in a pilot experiment in the SJLxB10.PL mouse model for EAE induced by MBP_Ac1-9_-peptide but developing relapses relying on PLP-specific T cells. We applied repeatedly pPLP_139_-_154_-PEG20 during ongoing EAE, but only a short delay in the first relapse was observed. However, in the remission phases, disease activity in this model does not completely disappear. Moreover, almost all mouse EAE models incorporate the application of CFA for induction. According to our above mentioned findings, the mycobacterial LPS from CFA is potentially contributing to the blockade of a tolerogenic effect. But also the entire concert of proinflammatory signals generated upon disease initiation might act by preventing the tolerogenicity of either native or carrier-conjugated peptide.

While these findings suggest that tolerogenic vaccination with PEGylated myelin-peptides alone is not effective in ongoing disease, the data demonstrate that the application even under the condition of already generated effector cells is safe, as we did not observe adverse effects resulting in exaggerated disease symptoms upon single, repeated or high dose application during EAE in the MOG or the SJLxB10.PL MBP/PLP mouse model. Also in the OVA-model, pOVA-PEG20 did not lead to an activated immune response in mice adoptively transferred with Th1 cells, to mimic an established disease. While both pOVA and pOVA-PEG20 induced proliferation of T cells, only the latter did not give rise to increased frequencies and numbers of IFN-γ-producing Th1 cells (Pfeil et al., Submitted). These findings might be of note with respect to reports on exacerbation of disease with myelin-derived native peptides (Genain et al., 1996; Bielekova et al., 2000; Kappos et al., 2000; Smith et al., 2005)

In the last decade, a variety of concepts to improve the efficacy of peptide tolerization has been developed. Apart from the loading of apoptotic cells with peptides already tested in a clinical trial (Lutterotti), different synthetic nanoparticles or liposomes have been loaded with antigenic tolerogenic self (HSP-) peptides, partly under additional encapsulation of tolerogenic adjuvants or modified surfaces to target the particle to specific compartments (Keijzer et al., 2013; Kishimoto and Maldonado, 2018; Pearson et al., 2019). In contrast to our findings with PEGylated peptides, the group of S. D. Miller could achieve resolution of ongoing EAE under therapeutic conditions with peptide coupled to polystyrene or biodegradable poly(lactide-co-glycolide) microparticles of distinct size and surface charge (Getts et al., 2012; McCarthy et al., 2017). The data suggest that these types of particles target specific subsets of APC and even without antigen actively contribute to stimulation of inhibitory pathways and tolerogenicity. Promising preclinical results in therapeutic settings were also obtained in the group of T. K. Kishimoto by combination of antigenic peptides with tolerogenic adjuvants such as rapamycin, most elegantly by loading nanoparticles with both, peptide and rapamycin (LaMothe et al., 2018).

## Conclusions

The present study has demonstrated that vaccination with PEG-coupled myelin peptide is superior to native peptide in inducing specific tolerance in the EAE model and acts predominantly by induction or expansion of Tregs. Straightforward chemistry and broad clinical experience with PEGylated biologicals are in favor of this concept. However, the weak effects of these agents on suppressing an already activated immune system are not sufficient to achieve therapeutic effects in ongoing disease. Whether subtypes of human MS with long and complete remission phases are potential targets of the concept remains to be shown, as we have not seen adverse activation of effector T cells by the PEG conjugates. A few other applications appear conceivable, e.g. for the inhibition of antibody formation towards intentionally applied foreign antigens, or for a safer variant of SIT to treat allergies, where so far mostly unmodified allergens are used. Moreover, PEG-conjugates might find a place in combination therapies with agents or treatments suppressing effector mechanisms. The anti CD3 therapy already being in clinical testing is such a candidate, as it appears to cause the deletion of effector/memory cells (You and Chatenoud, 2018). Other options would include the co-administration of immunosuppressants such as mAbs to costimulatory molecules and cytokines, or rapamycin.

## 6 Author Contributions

JP, MS, UH, RK, FL and AH conceived and designed the experiments. BT and RK synthesized and analyzed reagents, JP, MS, UL, CP, JPa and MK performed the experiments. JP, MS and PD analyzed the data. JP and AH wrote and edited the manuscript.

## 8 Acknowledgments

We would like to thank the labmanagers in the DRFZ for technical support, Friederike Ebner (Free University Berlin) for the introduction into the EAE model, Ping Shen (DRFZ) for technical advice in spinal cord isolation, Carmen Infante-Duarte (Charité) for critical reading and Frank Konietschke (Charité) for statistical advice.

## Supplementary Material

***Supplementary Method:*** The SJLxB10.PL model.

### Mice

SJLxB10.PL F1 mice were crossed in the breeding facility of the Deutsches Rheuma-Forschungszentrum Berlin (DRFZ). Mice were maintained under specific pathogen-free conditions according to national and institutional guidelines. All experiments were approved by the Landesamt für Gesundheit und Soziales (LAGeSo).

### Peptides

The peptides MBP_Ac1-9_ (Ac-ASQKRPSQR) and pPLP_139-154_ (HCLGKWLGHPDKFVGI) were synthesized in house (Institute for Medical Immunology, Charité Universitätsmedizin Berlin, Germany).

### Synthesis of pPLP_139-154_-PEG20

PEG20 was coupled to C_140_ of pPLP139-154. All other procedures were as described for pMOG-PEG20 in the main text.

### Induction of relapsing-remitting EAE in SJLxB10.PL mice and treatment protocol

RR**-**EAE was induced according to Miller et al., 2010 by *s*.*c*. immunization of SJLxB10.PL mice with 100 µg MBP_Ac1-9_-peptide in CFA (Difco, Heidelberg, Germany) supplemented with *Mycobacterium tuberculosis* 0.8 mg H37Ra on day 0. In addition, mice received 200 ng *Bordetella pertussis* (List Biological Laboratories, Campbell, USA) *i*.*p*. on day 0 and day 2, respectively. Mice were repetitively treated with PBS (control) or 50 µg (based on peptide amount) pPLP-PEG20 weekly from day 13.

## Supplementary Figures

**Supplementary Figure S1.**
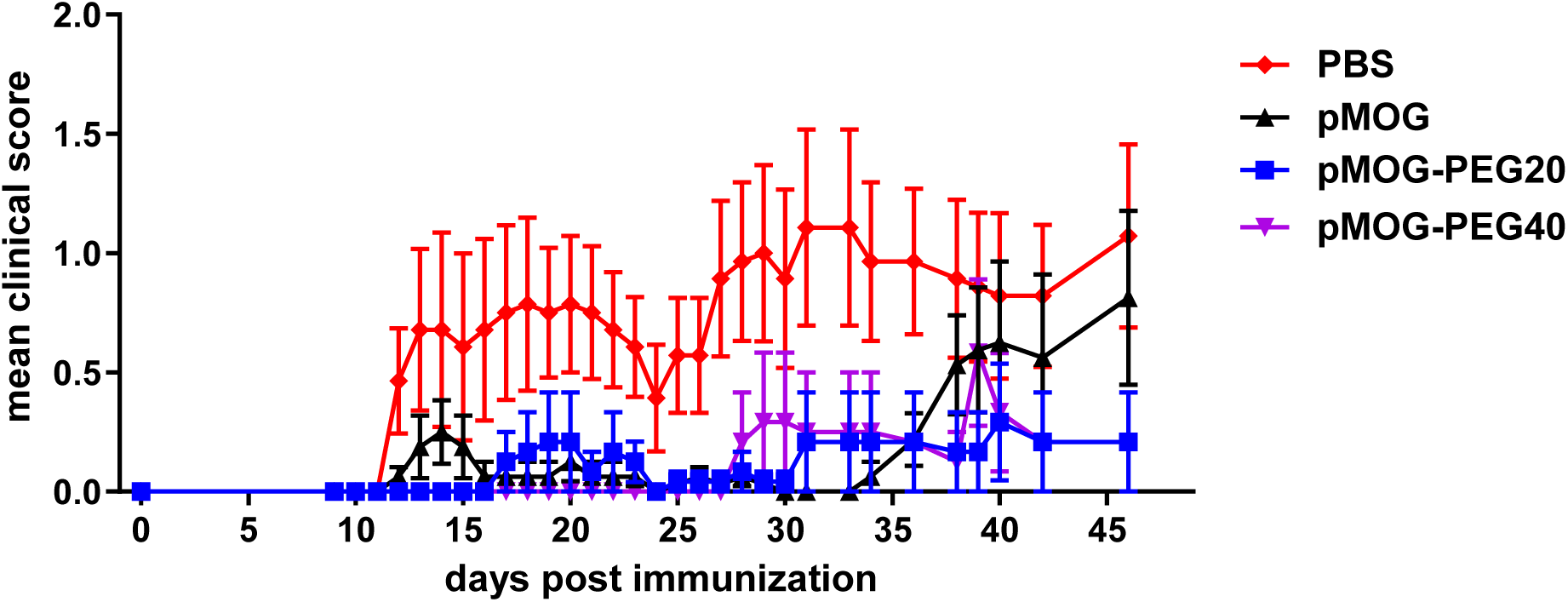
pMOG-PEG40 also ameliorates EAE symptoms. 14 days prior to EAE induction mice were tolerized with PBS (control), 7.6 µg pMOG, equimolar amounts of pMOG-PEG20 or equimolar amounts of pMOG-PEG40. Mean clinical score per group ± SEM is shown (n = 6-8 per group). One representative of two independent experiments is shown.

**Supplementary Figure S2.**
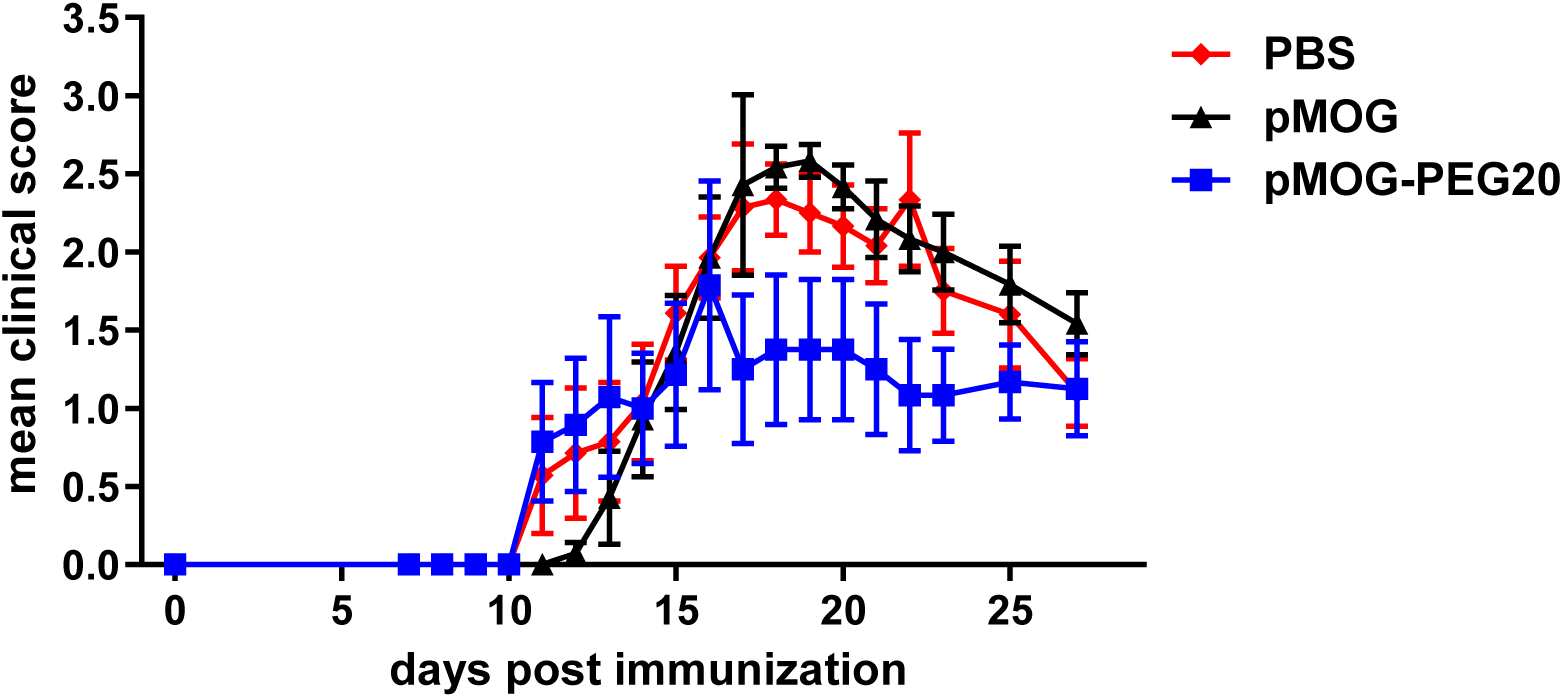
Reduced protection by pMOG-PEG20 upon administration four weeks prior to EAE induction. 28 days prior to EAE induction, C57BL/6 mice were tolerized *i*.*v*. with PBS (control), 7.6 µg pMOG or equimolar amounts of pMOG-PEG20. Mean clinical score per group ± SEM is shown (n = 7). One representative of two independent experiments is shown.

**Supplementary Figure S3.**
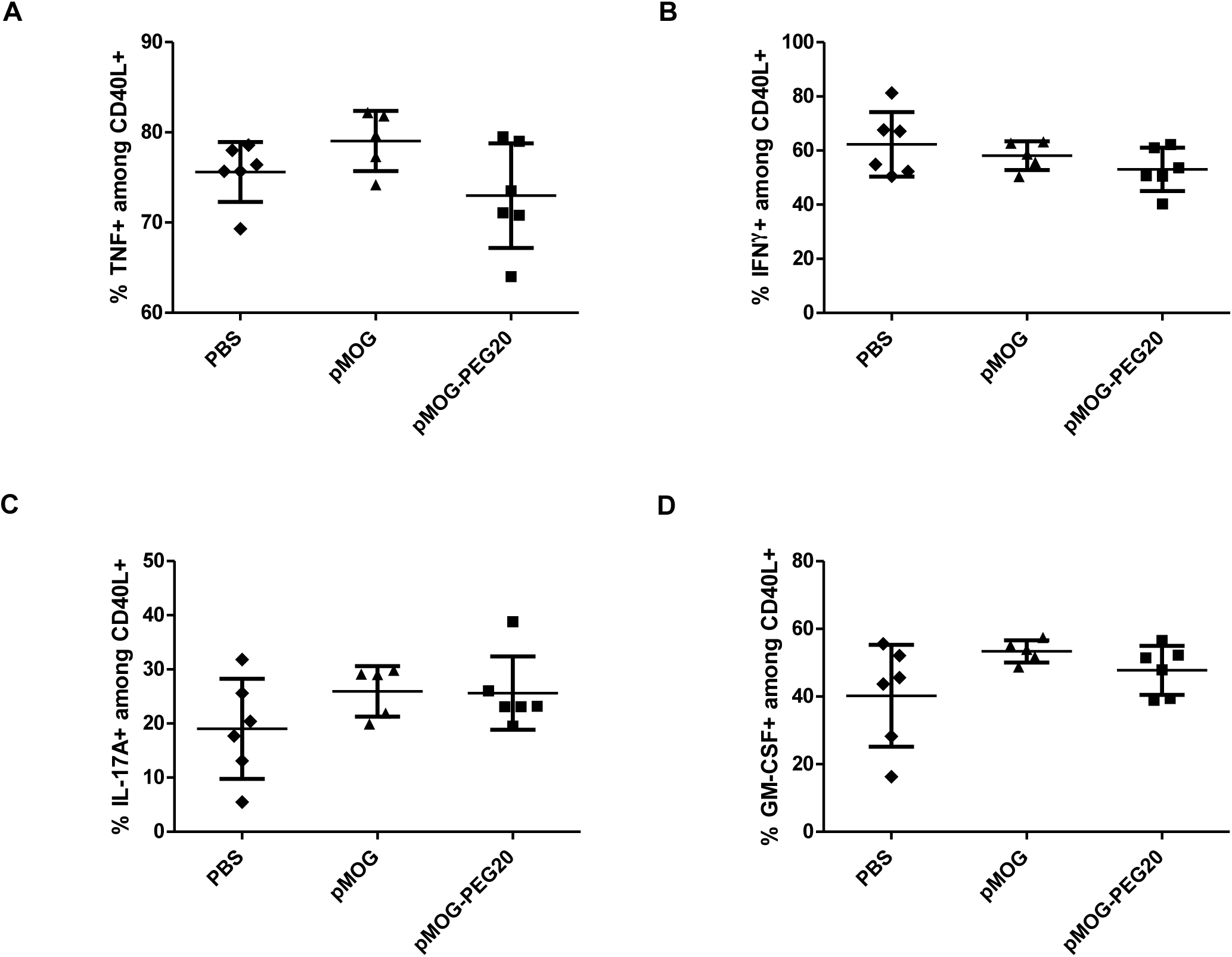
At the peak of disease, frequencies of proinflammatory cytokine producing CD4^+^ cells are not affected by pMOG-PEG20. C57BL/6 mice were tolerized *i*.*v*. with PBS (control), 7.6 µg pMOG or an equimolar amount of pMOG-PEG20 seven days prior to EAE induction. Animals were sacrificed at the peak of disease (d15 post EAE induction). Splenocytes were re-stimulated overnight with pMOG and analyzed by flow cytometry. Data represent mean of (**A**) % TNF^+^, (**B**) % IFN-γ^+^, (**C**) % IL-17A^+^ and (**D**) % GM-CSF^+^ splenocytes among CD4^+^ CD40L^+^ (MOG-specific) cells ± SD (n = 5–6 per group). One representative of two independent experiments is shown.

**Supplementary Figure S4.**
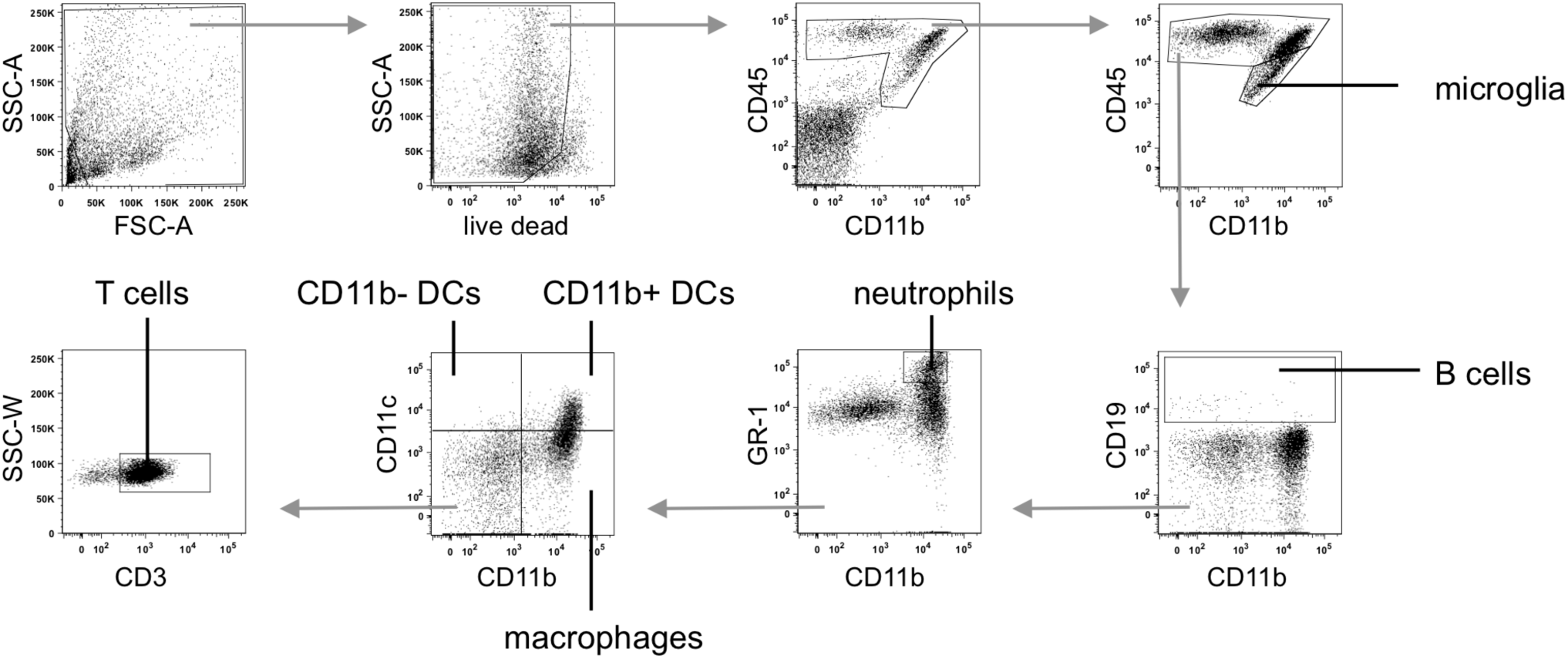
Sequential gating strategy used for the identification of leukocyte subsets in the CNS: microglia (CD45^int^ CD11b^+^), neutrophils (CD45^high^ GR1^high^ CD19^-^), CD11b^-^ DCs (CD45^high^ GR1^-^ CD11c^+^ CD11b^-^ CD19^-^), CD11b^+^ DCs (CD45^high^ GR1^-^ CD11c^+^ CD11b^+^ CD19^-^), macrophages (CD45^high^ GR1^-^ CD11c^-^ CD11b^+^ CD19^-^), B cells (CD45^high^ CD19^+^), and T cells (CD45^high^ GR1^-^ CD11c^-^ CD11b^-^ CD19^-^ CD3^+^).

**Supplementary Figure and Table S5.**
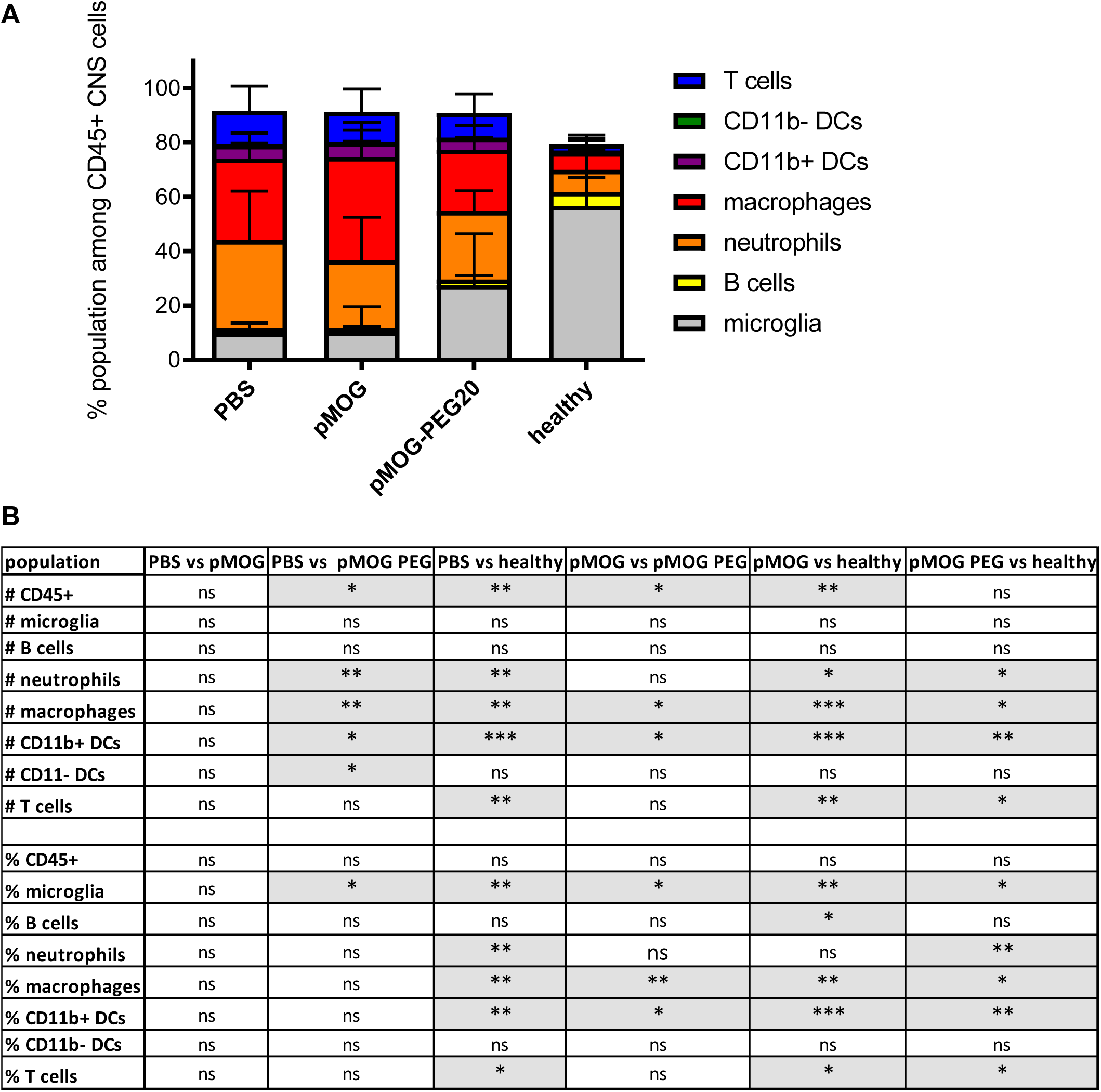
These data complement those of Figure 5 of main text. **(A)** Relative proportion of the analyzed subsets within CD45^+^ cells isolated from the CNS. Frequencies at the peak of the disease (d15) are depicted as mean ± SD (n = 6-12). Data is summarized from two independent experiments. **(B)** Table of statistical analyses. P values were determined by unpaired non-parametric Mann-Whitney U test.

**Supplementary Figure S6.**
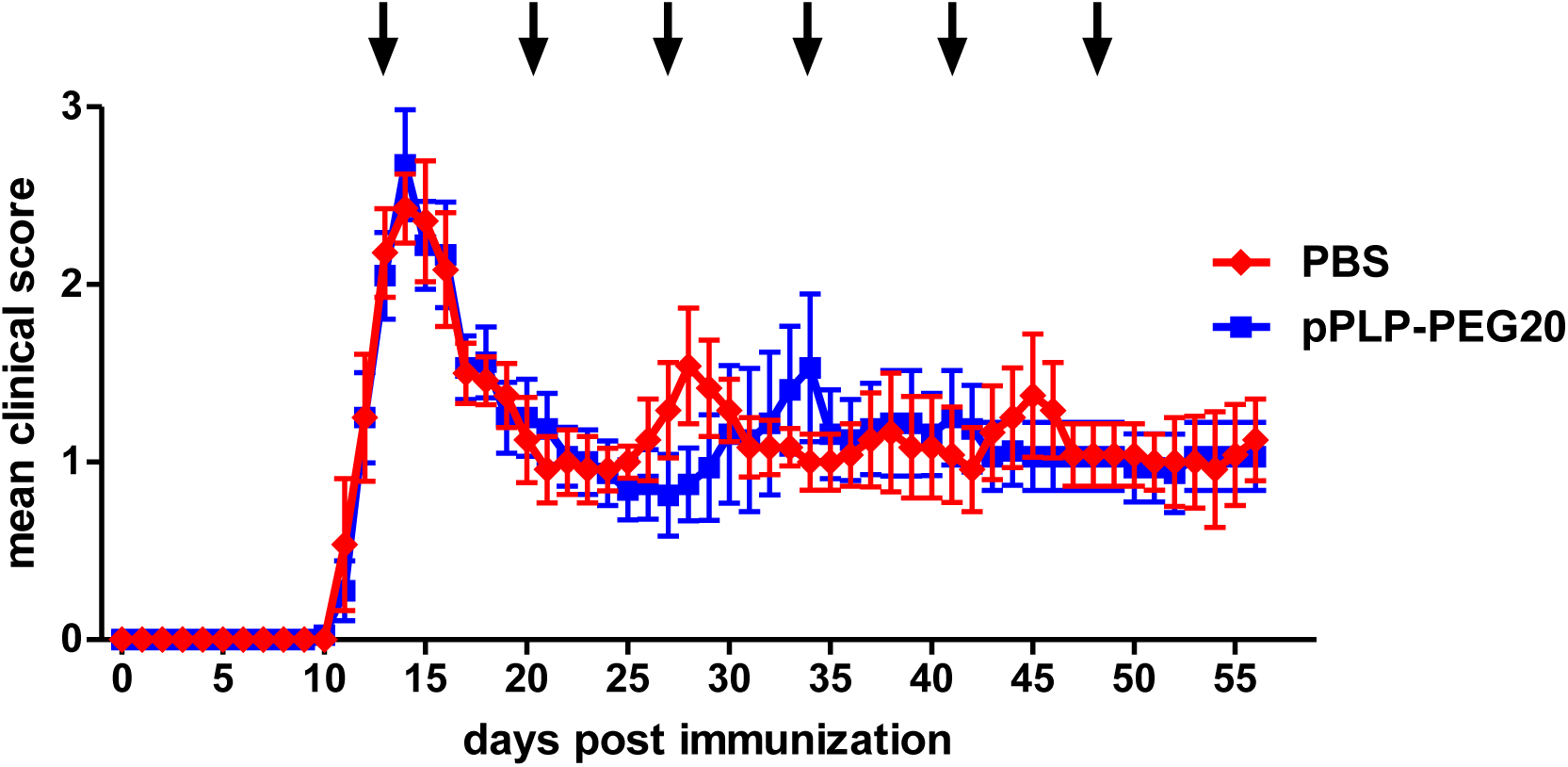
Repetitive treatment with pPLP_139-154_-PEG20 slightly delays, but does not prevent the pPLP_139-154_-specific relapse. RR**-**EAE was induced by *s*.*c*. immunization of SJLxB10.PL mice with MBP_Ac1-9_-peptide in CFA on day 0. In addition, mice received *Bordetella pertussis i*.*p*. on day 0 and day 2, respectively. Mice were repetitively treated with PBS (control) or 50 µg (based on peptide amount) pPLP-PEG20 weekly from day 13 (arrows). Mean clinical score per group ± SEM (PBS n = 7; pPLP-PEG20 n = 10). Data shown are from one experiment.

